# Insights into global planktonic diatom diversity: Comparisons between phylogenetically meaningful units that account for time

**DOI:** 10.1101/167809

**Authors:** Teofil Nakov, Jeremy M. Beaulieu, Andrew J. Alverson

## Abstract

Metabarcoding has offered unprecedented insights into microbial diversity. In many studies, short DNA sequences are binned into consecutively higher Linnaean ranks, and ranked groups (e.g., genera) are the units of biodiversity analyses. These analyses assume that Linnaean ranks are biologically meaningful and that identically ranked groups are comparable. We used a meta-barcode dataset for marine planktonic diatoms to illustrate the limits of this approach. We found that the 20 most abundant marine planktonic diatom genera ranged in age from 4 to 134 million years, indicating the non-equivalence of genera because some had more time to diversify than others. Still, species richness was only weakly correlated with genus age, highlighting variation in rates of speciation and/or extinction. Taxonomic classifications often do not reflect phylogeny, so genus-level analyses can include phylogenetically nested genera, further confounding rank-based analyses. These results underscore the indispensable role of phylogeny in understanding patterns of microbial diversity.

---

With potentially millions of species occupying all the world’s aquatic and terrestrial biomes, microbial species diversity is notoriously difficult to discover and catalog. Traditional approaches to species discovery are time and labor intensive, and they miss species that cannot be cultivated in the lab [1]. The phylogenetic diversity of this undiscovered “microbial dark matter” is often characterized through community DNA sequencing of barcode genes. A typical workflow includes DNA extraction from an environmental sample, PCR amplification of a barcode fragment, and high-throughput DNA sequencing of the amplicon. Sequencing reads are clustered into operational taxonomic units (OTUs) that are subsequently binned into consecutively higher taxonomic ranks, and these ranked groups, in turn, are often the focus of biodiversity assessments [2].

Linnaean names and ranks are often taken to mean more than what they are: arbitrary taxon delimitations disconnected from evolutionary history. The treatment of named ranks as anything other than arbitrary implies that identically ranked groups are somehow comparable, encouraging comparisons of their ecology, biogeography, and species richness [3, 4, 5]. The only meaningful comparisons involve groups with comparable evolutionary histories [6]. In this sense, monophyletic groups (clades) are more likely to be biologically cohesive units, and they should have comparable species richness if they are similar in age and have diversified at similar rates [7]. Comparison of monophyletic groups, while accounting for time, provides a robust framework for detecting clades with exceptional species richness and comparing their functional, ecological, or biogeographic breadth [8].

The Tara Oceans Project sequenced 18S-V9 metabarcode fragments from plankton samples to characterize microbial communities and species richness across the world’s oceans [9]. A total of 20 diatom genera accounted for nearly 99% of all diatom sequencing reads, and these genera were found to differ in relative abundance, cell size, habitat preference, geographical distribution, and species richness [2], however, it was unclear whether or not these patterns deviated from expectations.

We focused our analyses on the genus-based patterns of species richness and expected that older genera would be more species rich because they have had a longer period of time to diversify [7]. We used a time-calibrated phylogenetic tree of 1,151 diatoms [10] to calculate expected species richness for the 20 most abundant marine diatom genera in the Tara Oceans survey [2]. Given the crown age of diatoms [10], relative extinction (i.e., extinction/speciation) estimated from Cenozoic fossil species [11], and a minimum approximation of total described and undescribed diatom diversity (30,000 species; [12]), we calculated a net diversification rate for diatoms (i.e., speciation - extinction). We then used this rate to calculate the upper and lower bounds of expected OTU richness for the 20 focal diatom genera [8].

The 20 diatom genera ranged in age from 4134 million years (My). OTU richness was only weakly correlated with clade age (r=0.36, 95% CI=-0.1–0.7, df=18, P=0.12), with 12 of the 20 genera falling within expectations for OTU diversity given their age (Figure 1). The most abundant and OTU-rich genus, *Chaetoceros*, was also the oldest (Figure 1a). The birthdeath diversification model predicted that *Chaetoceros* diversity should range between 47 and 6567 species—the Tara Oceans dataset recovered 644 OTUs, consistent with expectations for a clade of this age (Figure 1b). Some of the most diverse genera identified by metabarcoding (e.g., *Corethron* and *Pseudo-nitzschia*) had OTU richness estimates that exceeded expectations. Assuming OTUs correspond to species and that our estimates of clade age are not heavily biased, these genera have either exceptionally high speciation or low extinction rates. Identifying the drivers of these patterns might offer new mechanistic insights into phytoplankton diversification. Comparisons between OTU richness (Figure 1b) and accepted taxonomic names from AlgaeBase (Figure 1c) showed expected discrepancies for lineages with substantial diversity in benthic or freshwater habitats (e.g., *Navicula*; Figure 1b, B and F annotations; Figure 1c, blue bars) and were also useful in highlighting clades that are underdescribed at the species level (Figure 1c, green bars).

**Figure 1:**
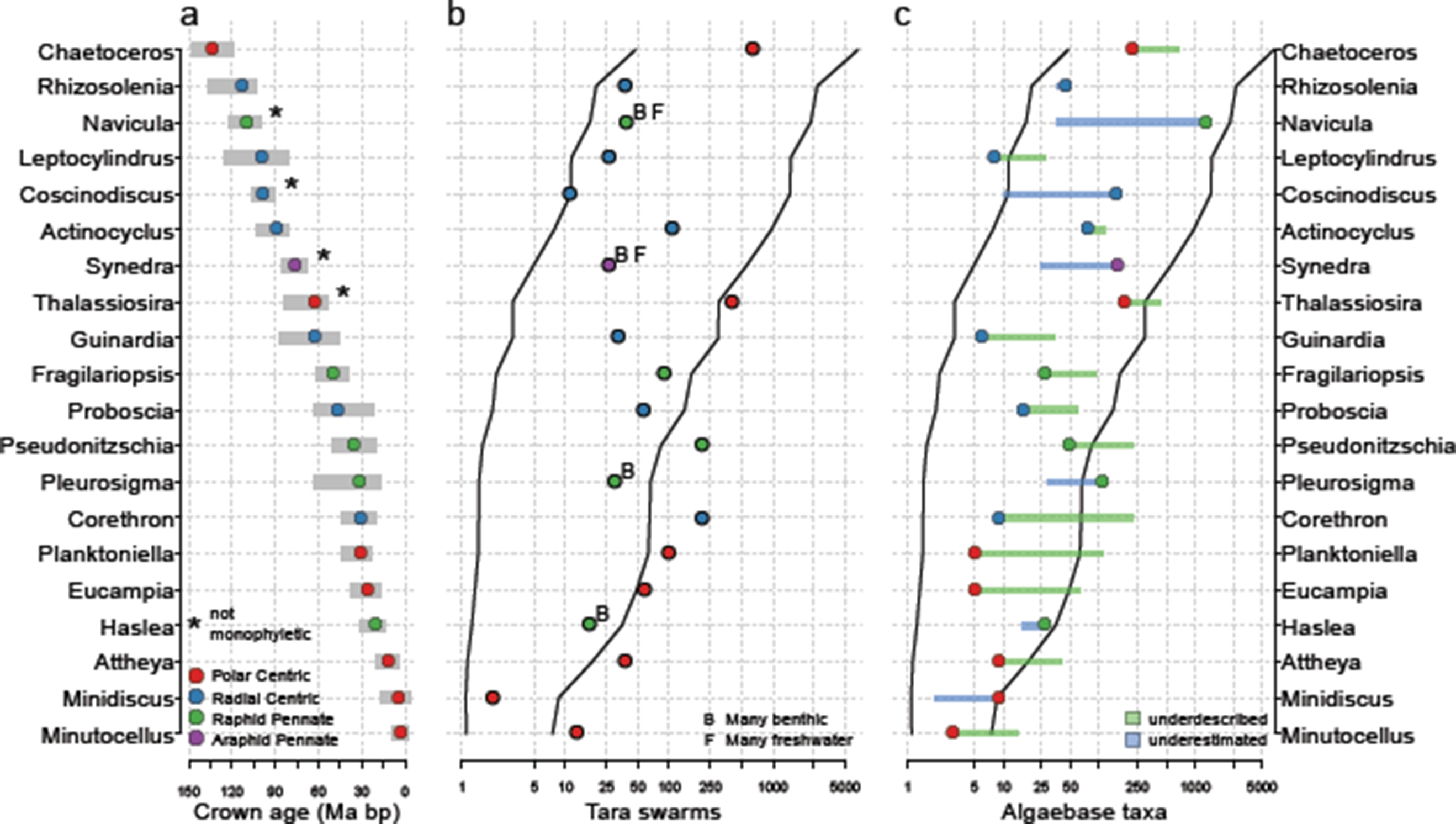
Age and estimated taxon richness of the 20 most abundant marine planktonic diatom genera identified by the Tara Oceans metabarcode project [2]. Crown ages and uncertainty (grey bars) were estimated from 1000 bootstrap phylogenies [10] (a). Taxon richness was estimated from the number of OTU swarms in the Tara Oceans dataset (b) and the number of accepted species names in AlgaeBase [13] (c); lines delimit 95% confidence intervals of expected richness given the crown age of a clade, empirical extinction fraction, and diatom-wide estimate of the net diversification rate.

Metabarcoding identified *Thalassiosira* as one of the most abundant, OTU-rich, and geographically widespread marine planktonic diatom genera. A total of eight Thalassiosirales genera were detected in the Tara Oceans project (*Cyclotella, Lauderia, Minidiscus, Planktoniella, Porosira, Shionodiscus, Skeletonema, and Thalassioisra*), and these genera ranged in age from 463 My (Figure 2). Thalassiosirales embodies many of the problems with misappropriation of biological or evolutionary properties to taxa based on their names [14]. The name *Thalassiosira* applies to a polyphyletic set of species whose common ancestor dates to 63 My and gave rise to nearly the full phylogenetic breadth Thalassiosirales diversity (Figure 2, diamond). As a result, including *Thalassiosira* in genus-level analyses leads to highly biased comparisons involving a genus that, in reality, corresponds roughly to a taxonomic order (Figure 2). Moreover, four of the eight thalassiosiroid genera detected by metabarcoding are nested within *Thalassiosira*, highlighting a common source of non-independence in rank-based comparisons (Figure 2, yellow branches). A more informative, phylogenetically based genus-level classification may have revealed clade-specific habitat preferences or geographic distributions among the many distinct *Thalassiosira* lineages [15].

**Figure 2:**
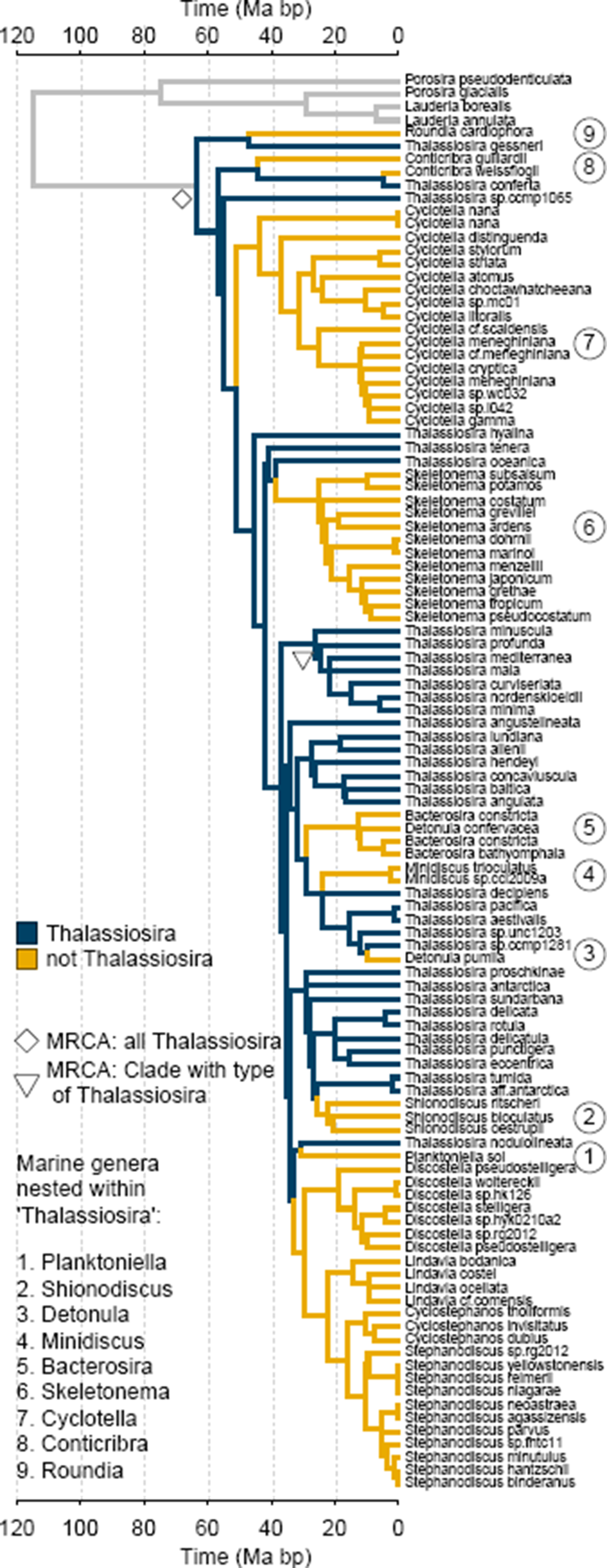
The genus Thalassiosira encompasses at least 10 marine planktonic diatom (including *Thalassiosira)* that range from 4-63 My in age. Topology and divergence times are based on [10].

The problems with rank-based comparisons, including as they relate to di-atoms, are well known [14, 16, 15]. A frequently cited advantage of metabar-coding is that it does not require taxonomic expertise. Still, the taxonomic affiliations of metabarcode sequences often become the units of biodiversity analyses. A working knowledge of phylogeny and systematics—which invariably highlight the deficiencies of Linnaean classifications—can lead to more meaningful analyses that explicitly incorporate phylogenetic history, ensuring robust comparisons of biologically equivalent units that account for time.

